# Designing Anti-Zika Virus Peptides Derived from Predicted Human-Zika Virus Protein-Protein Interactions

**DOI:** 10.1101/156695

**Authors:** Tom Kazmirchuk, Kevin Dick, Daniel. J. Burnside, Brad Barnes, Houman Moteshareie, Maryam Hajikarimlou, Katayoun Omidi, Duale Ahmed, Andrew Low, Clara Lettl, Mohsen Hooshyar, Andrew Schoenrock, Sylvain Pitre, Mohan Babu, Edana Cassol, Bahram Samanfar, Alex Wong, Frank Dehne, James. R. Green, Ashkan Golshani

## Abstract

The production of anti-Zika virus (ZIKV) therapeutics has become increasingly important as the propagation of the devastating virus continues largely unchecked. Notably, a causal relationship between ZIKV infection and neurodevelopmental abnormalities has been widely reported, yet a specific mechanism underlying impaired neurological development has not been identified. Here, we report on the design of several synthetic competitive inhibitory peptides against key pathogenic ZIKV proteins through the prediction of protein-protein interactions (PPIs). Often, PPIs between host and viral proteins are crucial for infection and pathogenesis, making them attractive targets for therapeutics. Using two complementary sequence-based PPI prediction tools, we first produced a comprehensive map of predicted human-ZIKV PPIs (involving 209 human protein candidates). We then designed several peptides intended to disrupt the corresponding host-pathogen interactions thereby acting as anti-ZIKV therapeutics. The data generated in this study constitute a foundational resource to aid in the multi-disciplinary effort to combat ZIKV infection, including the design of additional synthetic proteins.

## Introduction

The Zika virus (ZIKV) is currently causing an ongoing pandemic, incurring considerable human impact. The rapid spread of the virus throughout the Western hemisphere has driven a significant accumulation of knowledge on ZIKV infection [1–4]. The ZIKV is a positive-sense single stranded RNA (+ssRNA) arbovirus from the genus *Flavivirus* [5,6]. A mature ZIKV virion contains a monopartite segment of +ssRNA 10,800 nucleotides enclosed in a capsid comprised of C-proteins and surrounded by a 50 nm spherical envelope. The membrane is comprised of membrane (M) and envelope (E) proteins arranged around the icosahedral capsid. The entire genome is translated in to a single polyprotein 3,419 amino acids in length, which is cleaved at ten locations to produce 11 individual proteins. Despite significant similarity to other *Flaviviridae*, the ZIKV results in some symptoms that are not associated with members of this viral family such as Dengue or Yellow Fever [7].

Host-virus protein-protein interactions (PPIs) are essential for viral infection and propagation as well as neuroinvasion [8]. The ZIKV appears to be highly neuroinvasive (6.5X107 viral RNA copies/mg of brain tissue [9]) and has been linked to numerous neurological complications including congenital brain abnormalities [10], infant microcephaly [11], Guillain-Barré syndrome [12], and meningoencephalitis [13]. Additionally, the ZIKV has been found to cause testicular atrophy [14], and may be spread as a sexually transmitted infection [15].

Investigating the host-virus interactome is an important step in identifying targets for novel anti-viral therapeutics [16]. Designing molecules, such as competitive peptides that interfere with these PPIs, can serve as efficient anti-viral therapies [17–19]. An early example of such anti-viral peptides is S6 — a 111 amino acid long fragment of human integrase interactor protein 1 which forms a PPI with HIV-1 integrase protein. S6 is shown to be an effective inhibitor of HIV-1 replication [20].

Global PPI prediction analysis is a robust method for probing the network of host-pathogen interactions that occur at various stages of the viral life cycle. Recognition of pertinent PPIs between host cell proteins and ZIKV components can guide the development of synthetic inhibitory peptides capable of disrupting such interactions. In the current study, we use this approach to generate a list of specific peptide sequences that might function in combating ZIKV infection. We believe that the overproduction of these synthetic peptides may interfere with several human-ZIKV PPIs, thus interrupting the ZIKV lifecycle. Considering that the World Health Organization has lifted the declaration of emergency for the ZIKV, shifting to threat management of the ZIKV is important as many mechanisms of pathogenesis are still unclear. The designed peptides provided here may therefore prove to be useful therapeutics against ZIKV infection, and could aid in overall ZIKV management.

## Materials and Methods

### PIPE Prediction

From the suite of available sequence-based methods, the Protein-Protein Interaction Prediction Engine (PIPE) excels in terms of specificity and execution time [21]. PIPE was developed to investigate short co-occurring polypeptide sequences between two proteins to determine their likelihood of interaction [22–24]. This likelihood of interaction is captured by two scores: PIPE-Score and Similarity-Weighted score (further described in Pitre *et al*, 2008) [24]. Two datasets were used to perform the PIPE analysis, the first using the entire set of known human-virus interactions (irrespective of virus type) and the second considering only interactions specific to the *Flaviviridae, Herpeviridae*, *Arteriviridae*, and *Coronaviridae* families. PIPE analysis was applied to all combinations of human-ZIKV (20,515 human proteins, 11 ZIKV proteins). This resulted in a high confidence interaction network comprising the top-scoring 0.02% of predicted PPIs including 45 human-ZIKV interactions corresponding to 23 unique human proteins. From these 23 candidates, we chose the top 17 to further investigate based on their apparent relevance to human health.

### DeNovo Prediction

DeNovo is a host-virus PPI prediction tool tailored to predict cross-species protein interactions for newly identified viral organisms for which no interaction data are previously known [25]. Given the lack of known interactions for the ZIKV, DeNovo is well suited to this task. Similar to PIPE, DeNovo is a sequence-based method designed to leverage all currently known viral interaction data [25]. Unlike PIPE, however, DeNovo exploits the physiochemical properties of the human host’s proteins to inform its predictions [25]. Learning these characteristics in the commonly shared human host, across all known host-viral interactions, enhances the discovery of interactions in the organism of interest. Determining the high confidence interactions as those with a probability of interaction greater than or equal to 80% resulted in 871 human-ZIKV interactions (0.38% of all putative PPIs) corresponding to 186 candidate human proteins (supplementary table 1). From these 186 candidates, we performed a literature search on eight due to their relevance to ZIKV infection to supplement the 17 from the PIPE analysis.

**Table 1:** The 25 human protein candidates determined by PIPE (P) or DeNovo (N), respective ZIKV interactors, function (F), associated disease phenotype (D), and supporting literature.

| *Human Proteins* | *ZIKV Proteins* | *Function or Disease Phenotype* | *Reference* |
| --- | --- | --- | --- |
| ENO1 (P) | NS5, NS3, NS1*, M* | Affects cell proliferation, differentiation (F) | [29–31] |
| ENO2 (P) | NS5, NS3, NS1*, M* | Affects cell proliferation, differentiation (F) | [29–31] |
| ENO3 (P) | NS3, NS1*, E* | β-enolase deficiency (D) | [32] |
| RNASET2 (P) | NS3* | Microcephaly (D) | [33–36] |
| CALR3 (N) | NS4B, M | Male fertility, spermatogenesis, gamete fusion (F) | [37–39] |
| NME1 (P) | NS3 | Induction of fibronectin (F) | [40] |
| NME3 (P) | NS5, NS3, M | Cell motility in somatic cells, spermatozoa (D) | [41] |
| RNF151 (N) | Pr, M | Spermatogenesis (F) | [42] |
| RNF125 (P) | NS3 | Regulation of HIV-1 replication (F) | [43] |
| FUNDC1 (P) | NS3 | Activates hypoxia-induced mitophagy (F) | [44] |
| BCLG (P) | NS5, NS3, M | Apoptosis factor (F) | [45] |
| TRAF4 (P) | NS5, NS3, M* | Impaired neural crest development, folding (D) | [46] |
| MLPH (P) | NS3 | Griscelli syndrome (D) | [47] |
| PIAS3 (P) | NS3 | SUMOylation of photoreceptors (F) | [48] |
| CAMTA2 (P) | NS5, NS3, M*, C* | Decreased cardiac growth (D) | [49] |
| AZI2 (P) | NS5, NS3, M* | Neurodevelopment (D) | [50] |
| MATR3 (P) | NS3 | Cardiac development (F) | [51] |
| CEP63 (P) | NS5, NS3 | p53-dependent microcephaly (D) | [52] |
| RIAM (P) | NS5, NS3 | Leukocyte adhesion deficiency (D) | [53] |
| DYX1C1 (N) | NS4B, M | Dyslexia (D) | [54] |
| SNAP25 (N) | NS4B, M | Huntington's Disease (D) | [55] |
| YWHAE (N) | NS4B | Neurocognitive, Cerebrospinal fluid marker (D) | [56] |
| COX17 (N) | NS4B, NS2A | Cardiomyopathy, hepatic failure (D) | [57] |
| SPZ1 (N) | NS4B, M | Spermatogenesis (F) | [58] |
| DNAJA1 (N) | Pr, M | Testis development, spermatogenesis (F) | [59] |

\* indicates interaction pairs also predicted for the Dengue virus.

### Prediction of PPI-Sites

The list of PPIs generated from both methods can be used to inform the design of anti-ZIKV therapeutics by using peptide sequences from the predicted PPI site, which we refer to as the PPI-Site. We define the PPI-Site as the peptide sequence that is responsible for mediating the PPIs. When PIPE predicts an interaction between the two corresponding proteins, it will also report the predicted site of interaction using the amino acid sequences from both proteins. Using our previously reported PIPE output, we selected for PIPE’s built-in peak height attribute, which we refer to as H. This is a measure of the number of times two corresponding sequence pairs co-occur within the annotated database of known PPI normalized to its expected occurrence. Selecting for H within our top 17 interactions produced 4 human-ZIKV PPI-Sites which we believe mediate the human-ZIKV PPIs.

## Results

To generate a list of anti-ZIKV peptides, we first predicted a network of PPIs between humans and the ZIKV. This was accomplished using two complementary computational modeling methods — each trained on currently known high-confidence human-viral interactions obtained from VirusMentha (incorporating the MINT, IntAct, DIP, MatrixDB, and BioGRID databases) [26]. The first method, PIPE, is proficient in host-virus interaction prediction and has successfully been used to predict novel interactions in humans, yeast, and most recently in viruses (Hepatitis C and HIV-1) [22,24,27,28]. The second method, DeNovo, is specifically designed for the prediction of interactions between proteins from a host and newly discovered viral organisms independent of prior known host-virus interactions [25]. Figure 1 highlights the ontology classification of the 209 human proteins believed to participate in human-ZIKV PPIs based on their interaction profiles from both methods. In table 1, we report a total of 25 high priority human protein (and their associated ZIKV interactors) from our list of 209 candidates based on their correlation to known mechanisms of ZIKV infection, pathology, and symptomology. Of the eleven ZIKV proteins considered, the nine interacting proteins reported in our results correspond to the structural proteins: Envelope (E), Capsid (C), Premembrane (Pr), and Membrane (M), and the non-structural proteins: NS1, NS2A, NS3, NS4B, and the RNA-dependent RNA polymerase NS5 proteins. These proteins not only highlight protein candidates for PPI-based therapeutic design, but also they may provide potential mechanisms of ZIKV infection.

**Fig. 1.**
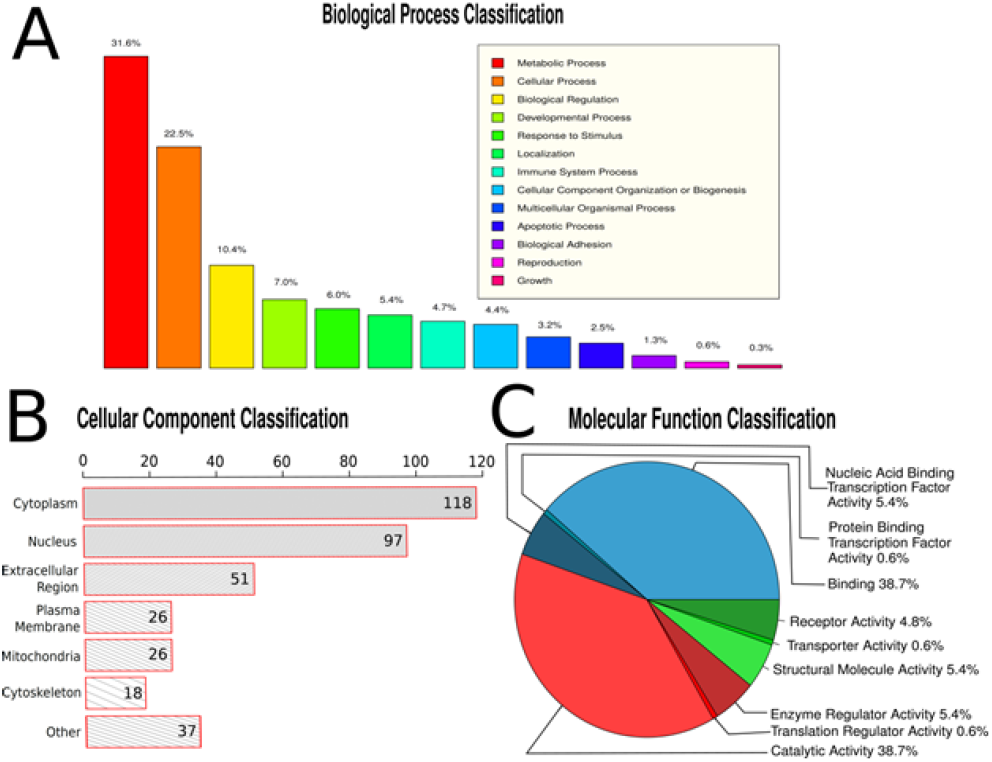
Human Protein Geneontology (GO) Classification using PANTHER GO-Slim. The evaluation of GO term enrichment for 198 of the 209 high confidence human proteins (8 unmapped proteins were excluded) for biological process (a), cellular component (b) and molecular function (c). Our list of 209 predicted high confidence human proteins can be found in supplementary table 1

From the list of 25 candidates, we here examine a subset of four proteins - RNASET2, ENO2, TRAF4, and CEP63 of particular interest for anti-ZIKV therapeutics. The first candidate, RNASET2, has been implicated in the occurrence of microcephaly [33] which is of special interest as no proposed mechanism exists linking the ZIKV to microcephaly. The second candidate, ENO2, has been linked to early brain development of humans [60], and may be useful in the study the pathogenesis of possible ZIKV-associated neurological disorders. TRAF4 is our third candidate and has been reported to be involved in TNF-receptor activity [46]. We believe that interfering with this receptor may affect ZIKV absorption and/or release. Finally, CEP63 has been previously implicated in p53 dependent microcephaly [52]. Considering that microcephaly is a hallmark of ZIKV infection in infants [61], designing a therapeutic which interrupts a ZIKV interaction with CEP63 may prove useful. We identified the site of interactions (PPI-Site) between these proteins and their ZIKV interacting partners (table 2). Overproduction of these PPI-Sites via synthetic peptides can interfere with human-ZIKV PPIs, potentially acting as competitive inhibitors against the corresponding ZIKV protein. Consequently, PPI-Sites can provide the basis for effective anti-ZIKV therapeutics.

**Table 2:**
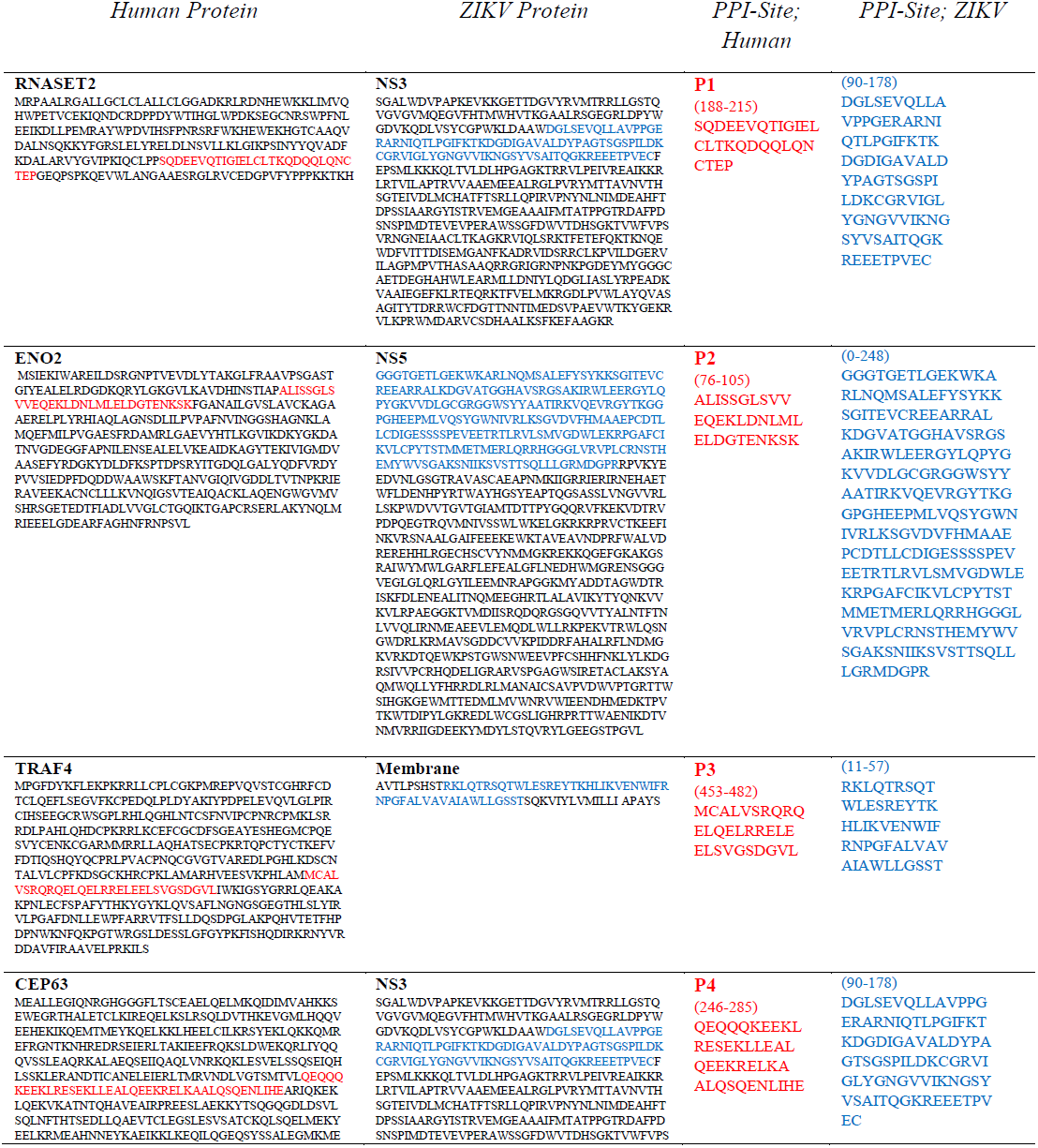
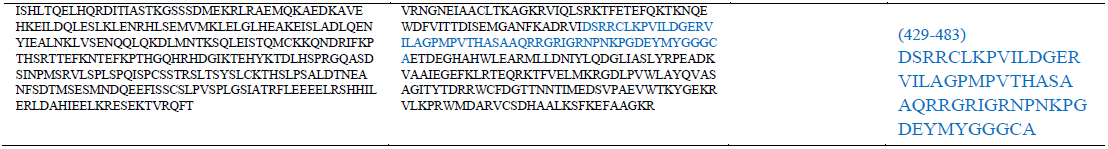
The amino acid sequence for Human-ZIKV PPIs. The peptides believed to mediate the interactions are indicated. P1-P4 represent peptides that may function as anti-ZIKV therapeutics. These peptides may interact with the corresponding ZIKV proteins and prevent them from interacting with their human PPI partners.

## Discussion

At the outset of this study, we sought to design anti-ZIKV therapeutics by identifying PPI-Sites from predicted human-ZIKV PPIs. By identifying these PPI-Sites on both the host and viral proteins, we provide a scaffold for the design of peptides which may inhibit ZIKV proteins or compete for binding to functionally-relevant targets. We employed two computational tools to predict PPIs, namely PIPE and DeNovo, and produced a priority list of testable protein interaction candidates which can be used as an important resource for multiple disciplines. In particular, the amino acid sequences identified here as being responsible for mediating interactions between human and ZIKV proteins can serve as the basis for engineering additional anti-ZIKV therapeutics.

One of the primary public health concerns of the ZIKV infection has been occurrence of microcephaly in newborn children who are infected with the virus, or whose mothers have been infected. It is therefore possible that the virus is acting on the fetus during pregnancy, and affecting development. Our candidate, RNASET2 has been causally linked to a range of neurologic impairments including microcephaly, multifocal white matter lesions, and anterior temporal lobe subcortical cysts [33]. RNASET2 has been shown to localize to the lysosome, where it may function in RNA catabolism [62]. Zebrafish models have established that the loss of function in RNASET2 results in neuronal lysosomal disorder [63] congruent with findings in humans that have linked lysosomal disorders to the development of microcephaly [34]. Moreover, a loss of function mutation at amino acid 184 in the RNASET2 protein has been found in infants with cystic leukoencephalopathy leading to microcephaly, white matter lesions, and other temporal lobe deficiencies [33].

We predicted an interaction between the human protein RNASET2 and the ZIKV serine protease NS3. Interestingly, two *Flaviviridae* (Hepatitis C and Dengue) have been found to interact with human RNASET2 protein via their NS3 protein *in vitro* [64]. The proteolytic domain of the ZIKV NS3 serine protease occurs at residues 1-175 [65,66], and RNASET2 is a proteolytic target at residue 24-25, which when cleaved produces a signaling peptide and the functional domain [67]. Within this functional domain exists a serine at residue 188, congruent with our predicted site of interaction (PPI-Site, residue 188-215) and four residues downstream of the leukoencephalopathy-associated mutation. Interestingly, we predict that this site interacts with the proteolytic domain of NS3 (PPI-Site, residue 1-178) which may impact the functionality of RNASET2, as the functional domain would theoretically be cleaved in two (Figure 2). Disrupting this interaction via introduction of the identified PPI-Sites (P1) in excess could disrupt the ZIKV lifecycle in the cell. Alternatively, the binding of the PPI-Site to the proteolytic domain of NS3 can interfere with the mode of action of NS3 by deactivating its proteolytic domain in a PPI independent manner. In both cases, excess of P1 may interfere with ZIKV infection and hence may function as an effective anti-ZIKV therapy.

**Fig. 2.**
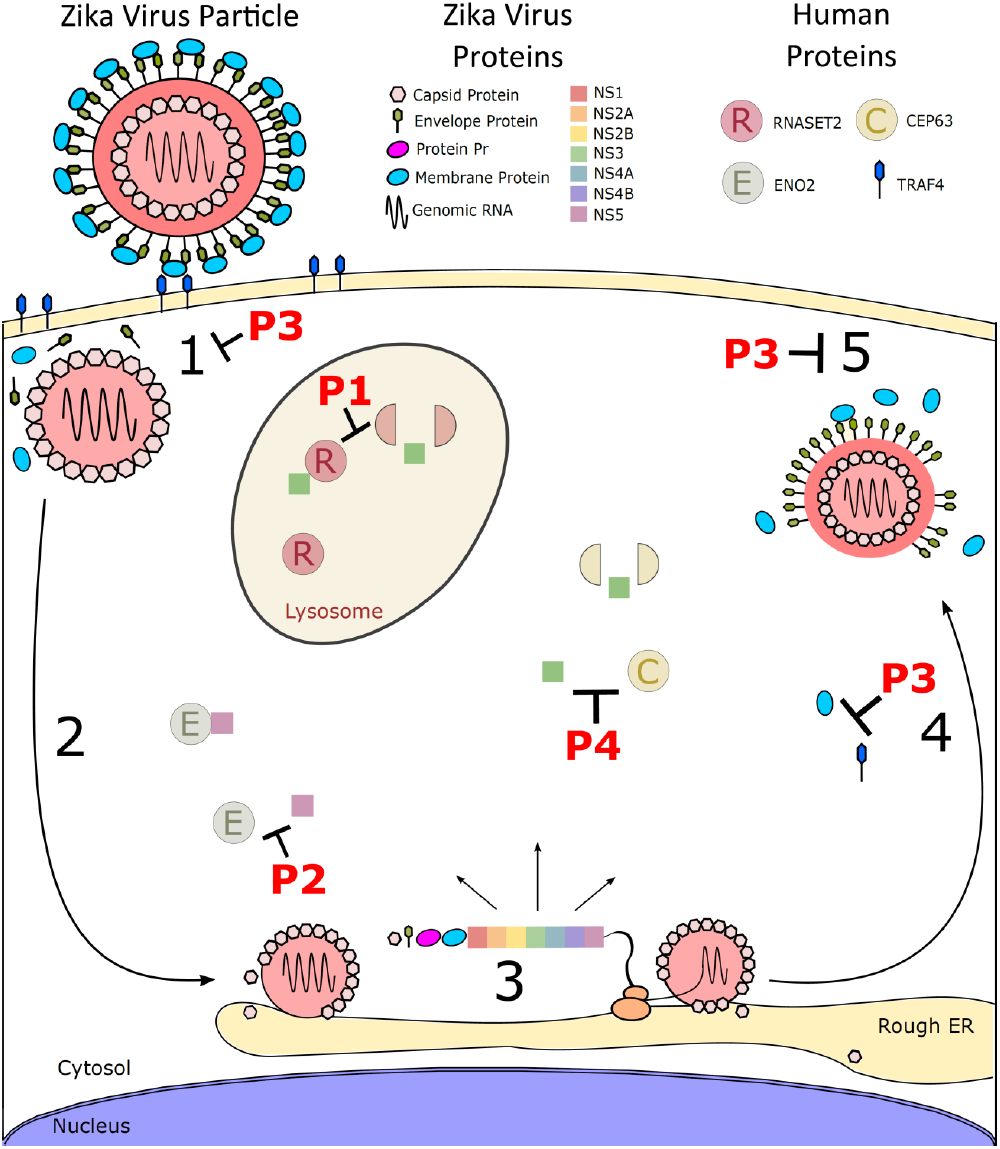
The ZIKV lifecycle within a human cell, and the peptide drugs that may interfere with its different steps. 1: The ZIKV particle adsorbs into the human cell, possibly via TRAF4. P3 may interfere with this step. 2: The virion is uncoated in the cytoplasm. 3: ZIKV RNA undergoes translation to produce the ZIKV polypeptide. P1, P2 and P4 are designed to interfere with the activity of synthesized ZIKV polypeptides. 4: The immature virion is assembled and moves to the cell membrane. 5: The ZIKV particle exits the host cell. P3 may also interfere with these last two steps. The ENO2 interaction with NS5 may impact neurodevelopment. This is the proposed site of P2 activity. The predicted interaction between RNASET2 and NS3, which we hypothesize to cleave the RNASET2 functional protein at amino acid 188 is the predicted site of P1 activity. The interaction between CEP63 and NS3 may result in a cleaved CEP63 protein representing the proposed site of P4 activity. P1: SQDEEVQTIGIELCLTKQDQQLQNCTEP, P2: ALISSGLSVVEQEKLDNLMLELDGTENKSK, P3: MCALVSRQRQELQELRRELEELSVGSDGVL, and P4: QEQQQKEEKLRESEKLLEALQEEKRELKAALQSQENLIHE.

Our additional peptide sequences come from the interactions between ENO2-NS5, TRAF4-M, and CEP63-NS3, as seen in Figure 2. These three additional human protein candidates are all involved in neurodevelopment. Specifically, ENO2 (neuron-specific enolase) is important for normal brain development in humans [30]. A member of the enolase family, ENO2 promotes cell maturation in the central nervous system [68]. During early development, neuronal cells switch from ENO1 (non-neuronal enolase) to ENO2 in order to promote brain development [29] (Figure 2). Interestingly, a similar interaction was reported by Munoz *et al.* where they describe an interaction between the *Aedes aegypti* enolase and Dengue virus NS5 protein [69]. The predicted P2 peptide could potentially act as competitive inhibitor for the NS5 binding site. In this way, we predict that P2 will interrupt the lifecycle of the ZIKV in the human cell, thus acting as an effective therapeutic.

A previous study identified AXL as a receptor candidate that is active in the ZIKV lifecycle [70]. Although not a receptor, TRAF4 is an essential element for normal TNF receptor function as TRAFs have been previously identified as adaptor proteins for membrane receptors [71]. Therefore, it is possible that the ZIKV M protein may be impacting functionality of the TNF receptor via TRAF4, possibly increasing the function of TNF. Pertaining to neurodevelopment, it has been shown that TRAF4 is required for normal neural crest formation [46]. If the ZIKV M protein is binding to TRAF4 in a inhibitory manner, it is possible that TNF receptor functionality may be impacted [71], and that neural crest formation may be negatively affected [46]. The corresponding PPI-Site derived peptide for this interaction is P3. Overproducing this peptide may interfere with ZIKV adsorption/penetration and thus act as a promising anti-ZIKV therapeutic. As well, if the ZIKV M protein is inhibiting TRAF4 leading to impaired neural crest formation, inhibiting the M protein via P3 would be ideal.

Finally, a short motif within centrosomal protein CEP63 was predicted to interact with the ZIKV serine protease NS3. This is an important interaction as the loss of CEP63 in mice results in microcephaly [52]. It is therefore possible that the NS3 serine protease could act on one of the two serines within the interaction site on CEP63, rendering the protein dysfunctional. Overproduction of P4 may be able to saturate NS3 activity and prevent detrimental proteolytic cleavage of the endogenous target.

In this study, we have identified proteins and PPI-Sites of interest and designed a series of peptides that could be used to combat ZIKV infection. These peptides theoretically interfere with the activity of their corresponding ZIKV proteins by competing for their interacting partners. Additional manipulation and/or modification of the peptides designed in this study can lead to new anti-viral therapeutics with improved properties such as increased half-life or stronger binding properties. The information provided here can be used to develop and engineer viral therapeutics for the ZIKV, a process that has been achieved with other viruses and is proposed for the ZIKV [72,73].

## Acknowledgments

This work was funded by the Natural Sciences and Engineering Research Council of Canada (NSERC). The authors claim no conflict of interest.

